# Left-right leaf asymmetry in decussate and distichous phyllotactic systems

**DOI:** 10.1101/043869

**Authors:** Ciera C. Martinez, Daniel H. Chitwood, Richard S. Smith, Neelima R. Sinha

## Abstract

Leaves in plants with spiral phyllotaxy exhibit directional asymmetries, such that all the leaves originating from a meristem of a particular chirality are similarly asymmetric relative to each other. Models of auxin flux capable of recapitulating spiral phyllotaxis predict handed auxin asymmetries in initiating leaf primordia with empirically verifiable effects on superficially bilaterally symmetric leaves. Here, we extend a similar analysis of leaf asymmetry to decussate and distichous phyllotaxy. We found that our simulation models of these two patterns predicted mirrored asymmetries in auxin distribution in leaf primordia pairs. To empirically verify the morphological consequences of asymmetric auxin distribution, we analyzed the morphology of a tomato *sister-of-pinformed1a* (*sopin1a*) mutant, *entire-2*, in which spiral phyllotaxy consistently transitions to a decussate state. Shifts in the displacement of leaflets on the left and right sides of *entire-2* leaf pairs mirror each other, corroborating predicted model results. We then analyze the shape of >800 common ivy (*Hedera helix*) and >3,000 grapevine (*Vitis* and *Ampelopsis* spp.) leaf pairs and find statistical enrichment of predicted mirrored asymmetries. Our results demonstrate that left-right auxin asymmetries in models of decussate and distichous phyllotaxy successfully predict mirrored asymmetric leaf morphologies in superficially symmetric leaves.

## INTRODUCTION

Leaf shape is defined across four axes. Asymmetry is usually obvious along three of these axes—the proximal-distal, abaxial-adaxial, and medio-lateral (Figure 1A), while asymmetry along a fourth axis, which describes the left and right halves of a leaf (Figure 1B), is rarely explored. Most leaves are thought to be either truly bilaterally symmetric (i.e., each side of the leaf mirrored along the midrib) or if asymmetry is observed, on average this asymmetry cancels out. Fluctuating asymmetry observed along the left/right axis of leaves has been attributed to noise created from random errors during development, influenced by environmental stresses [1–3]. Random noise in development can be assumed if the average difference between the left and right sides is zero, but if there is a consistent shift in asymmetry on one side compared to the other, directionalized asymmetry may be occurring [4].

**Figure 1:**
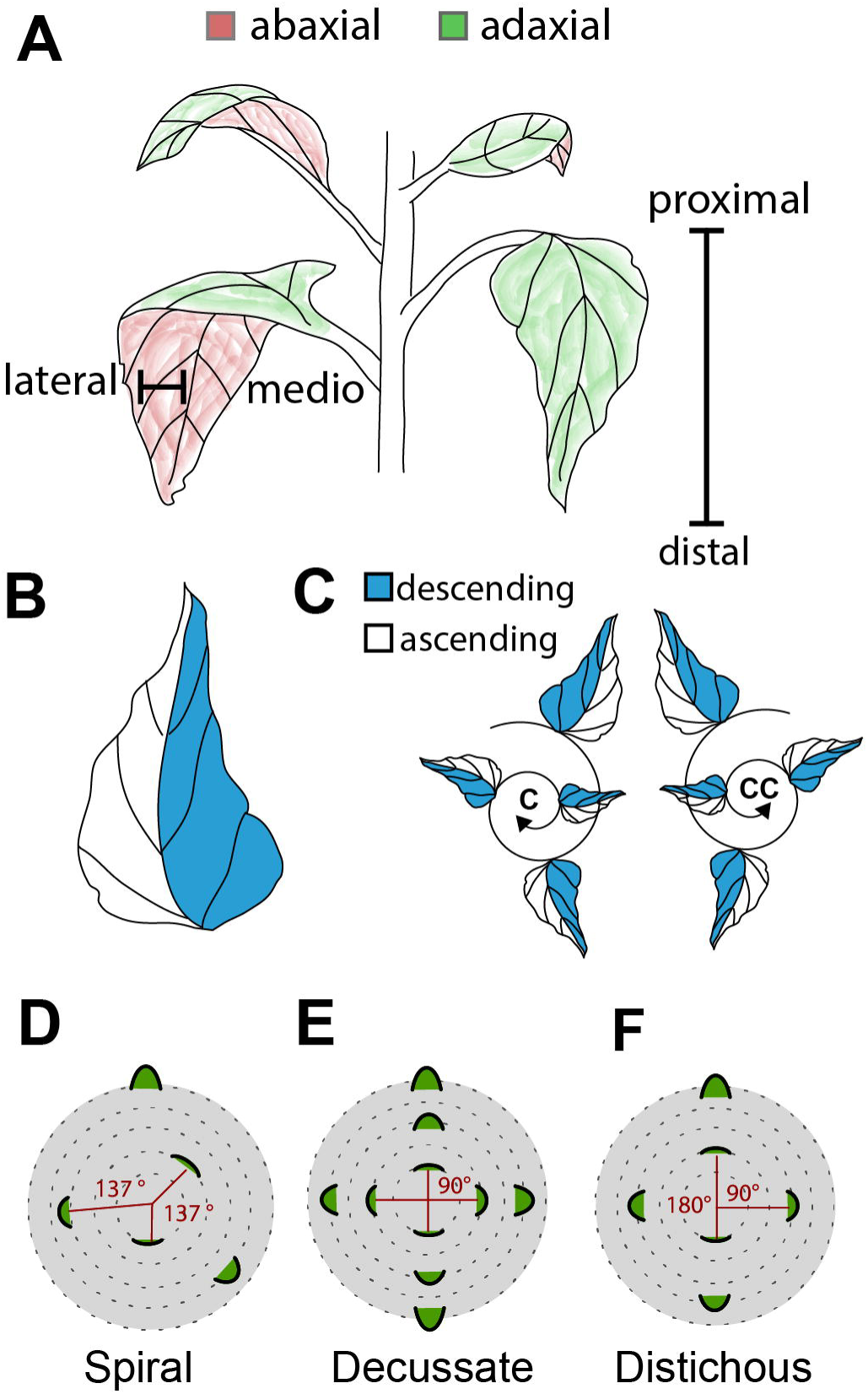
Leaf morphology is defined by four axes and the position of the leaf is defined by phyllotaxis. **A**) Schematic describing three developmental axes of a leaf. The medio-lateral axis describes the mid-vein to margin region of a leaf. The adaxial-abaxial axis defines the top and bottom of the leaf, respectively. The distal-proximal axis refers to the leaf tip relative to the base. **B-C**) The descending and ascending axis of a leaf refers to the left and right sides of the leaf in relation to spiral phyllotactic patterning. Spiral phyllotaxy can be further defined as Clockwise (C) or Counter-Clockwise (CC). The ascending side of the leaf faces the direction of younger leaves, while the descending side faces older leaves. **D**) When the divergence angle between each leaf is ~137.5° the phyllotaxis is defined as spiral. Decussate phyllotaxis refers to leaf initiation patterning where leaves initiate in pairs 180° apart, with 90° between each successive pair (literally “crossed”). **F**) Distichous phyllotaxis occurs when leaves initiate one at a time 180°apart (literally “of two rows”).

Recent work using sensitive techniques to measure leaf shape has suggested that directionalized asymmetry along the left and right side of the leaf may be more prevalent than once believed. Although superficially bilaterally symmetric, Elliptical Fourier Descriptor analysis in *Arabidopsis thaliana*, in addition to simple measurements in the shifts between leaflet placement along the left and right sides of *Solanum lycopersicum* (tomato) leaves, reveals that plants in these species produce leaves biased to be left‐ or right-handed [5]. The asymmetry of leaves is dependent on the handedness of the plant from which they originate. Handedness in plants arises when the phyllotaxy (the angular arrangement of initiated leaves and other lateral organs on a plant) is spiral (i.e., the angle between initiating leaves is approximately the golden angle, ~137.5°). In reference to a “bottom-up” view of leaf initiation events, the spiral can form in two directions, either “clockwise” (C) or “counter-clockwise” (CC) (Figure 1C). The observation that leaf morphological features are asymmetrically skewed in relation to spiral phyllotactic patterning has been described in exceptionally asymmetric species with respect to features such as venation patterning, leaf shape, curling, coiling, and resupination [6, 7]. Notably, many of the examples in the previous references are from the Zingiberales, including banana (*Musa*) and *Calathea*, reported to have a fixed phyllotactic handedness [7, 8]. The left and right sides of a leaf in plants with spiral phyllotaxy therefore appear to possess intrinsic asymmetries influenced by whether the phyllotaxy of the plant is clockwise or counter-clockwise (Figure 1C). To reflect the orientation of a leaf relative to the spiral phyllotactic direction, various terminologies have been proposed [7, 9–11]. In this paper, the leaf half which faces towards emerging leaves will be described as ascending, while the half facing the older leaves is the descending side (Figure 1C).

Modeling and experimental approaches have shown that phyllotactic patterning is largely determined by the localization of the plant hormone auxin [12, 13]. Regions of high auxin concentration determine sites of leaf initiation and the positional information of many other leaf features during morphogenesis, including vasculature, leaflets, and the development of the margin [14, 15]. The dynamics of auxin flux in spiral phyllotactic systems are such that it is predicted that auxin would be depleted from the ascending side of older primordia and supplied to the descending side of younger primordia. This prediction was verified both in existing auxin flux models of the shoot apical meristem (that have provided foundational insights into the self-organizing mechanisms by which auxin flux creates phyllotactic patterns) and empirically, by looking at the asymmetric shape features of initiating and mature leaves and the distribution of auxin reporter activity in leaf primordia [5]. While leaf asymmetry is guided by biased auxin distribution during spiral phyllotactic patterning, no one has explored how leaf asymmetries manifest in plants which display other phyllotactic patterns.

While the most common phyllotactic pattern in the plant kingdom is spiral, phyllotaxy can take on a multitude of orientations, including decussate and distichous. In decussate patterns pairs of leaves initiate at the same time 180° apart with subsequent leaf pares initiating at 90° from previous pair (Figure 1E). Distichous phyllotaxy is similar except leaves initiate one at a time, at a divergence angle of 180° (Figure 1F). In spiral systems the left-right axis is fixed and the developmental context of each side of a primordium depends on phyllotactic chirality, but in decussate and distichous phyllotaxy, the potential relationship between asymmetry in leaves relates to their initiation as pairs.

To explore the asymmetric nature of leaves arising from decussate and distichous systems, we asked if leaf asymmetries exist using both modeling and empirical approaches. We first revisited an auxin flux model that is capable of recapitulating decussate and distichous phyllotaxy. We found that there are predicted auxin asymmetries in initiating decussate and distichous leaf primordia pairs (that is, the divergence angle between the pair is ~180°) such that auxin falls on opposite sides of each leaf. Curious if such a mirrored relationship is empirically supported, we took advantage of a recently characterized *sister-of-pin-formed1a* (*sopin1a*) mutant in tomato, *entire-2* (*e-2*), that displays a consistent decussate phyllotaxy at early nodes [16]. Leaf pairs in *e-2* exhibit mirrored shifts in the displacement of leaflets on the left and right sides of the leaf, verifying modeled predictions. We then analyze the morphology of >800 distichous leaf pairs using Elliptical Fourier Descriptors in common ivy (*Hedera helix*) and find a statistical enrichment for successive leaves to have opposite asymmetry above that expected by chance. A similar analysis using homologous Procrustes-adjusted landmarks in >3,000 leaf pairs from 20 different species and hybrids of grapevine (*Vitis* and *Ampelopsis* spp.) shows that the predicted mirrored asymmetries in successive leaves are particularly strong in this group. Our results show that even in decussate and distichous species seemingly lacking directional asymmetries, models of asymmetric auxin flux successfully predict left-right asymmetries in mature leaves.

## RESULTS

### Modeling predicts mirrored IAA shifts between pairs of leaves in decussate and distichous phyllotaxy

Previous work asked if there are differences in auxin localization with respect to initiating leaf primordia using models capable of simulating spiral phyllotactic patterning [5]. We re-visit this model, which is capable of predicting many phyllotactic patterns, including spiral, decussate, and distichous [13]. The model works by simulating known mechanisms that direct leaf initiation events. The simulation reiterates through several steps. 1. Directional auxin transport by the action of the auxin efflux transporter, PIN-FORMED1 (PIN1), which directs auxin towards neighboring cells with the highest auxin concentration. 2. Auxin accumulates to convergence points. 3. Once auxin levels reach a certain threshold, auxin is transported inward beginning leaf initiation on the periphery of the simulated shoot apical meristem (SAM). Unlike the spiral and decussate models, in the distichous model the original equations from Smith and co-workers are stable enough to be used such that original transport equations, rather than those assuming no primordium differentiation, are used without *PIN* polarity bias in primordium cells. Thus our methods here are independent of the particular choice of transport equation, as several have been proposed [13, 17, 18].

The analysis of decussate and distichous systems is different from spiral phyllotaxis, as pairs of primordia are analyzed for relationships between their divergence angles and IAA Shift values. For the decussate system, a pair of leaves has to be preceded by a divergence angle between 70° and 100°, and the pair itself requires a divergence angle >150°. In the distichous system, a “pair” of leaves consists of two successive leaves in which the divergence angle is >130°. To facilitate analysis, divergence angles were converted to a positive sign that forces a similar orientation in all leaf primordia pairs. We then analyzed the relationships between divergence angles and IAA Shift values (the deviation of the center of mass of auxin distribution in a leaf primordium relative to the divergence angle) between “primordium A” and “primordium B” (see Figure 2A). If IAA Shift values between a pair of distichous or decussate primordia are of the same sign (e.g. shifting clockwise in both primordia, or counter-clockwise in both primordia) then we expect auxin maxima to fall on the same side of each primordium such that they are of a “pinwheel” orientation (Figure 2B). Contrastingly, if the IAA Shift values between primordium A and B are opposite in sign, the auxin maxima fall on mirroring sides of a primordia pair, forming a “mirror” orientation (Figure 2C). In both decussate and distichous models, we find that IAA Shifts form a mirroring pattern within primordia pairs (Figure 2D-F). In primordium A, as the divergence angle decreases from the ideal 180°, IAA Shift values decrease and become negative, whereas in primordium B as the divergence angle decreases, IAA Shift values increase and become significantly positive. The results show that as decussate or distichous pairs of leaves deviate from an ideal 180° divergence angle the auxin peak concentrations remain truer to 180°, creating mirror image pairs of primordia as in (Figure 2C) that is often empirically observed, as in distichous arrangements of *Begonia* leaves (Figure 2G).

**Figure 2:**
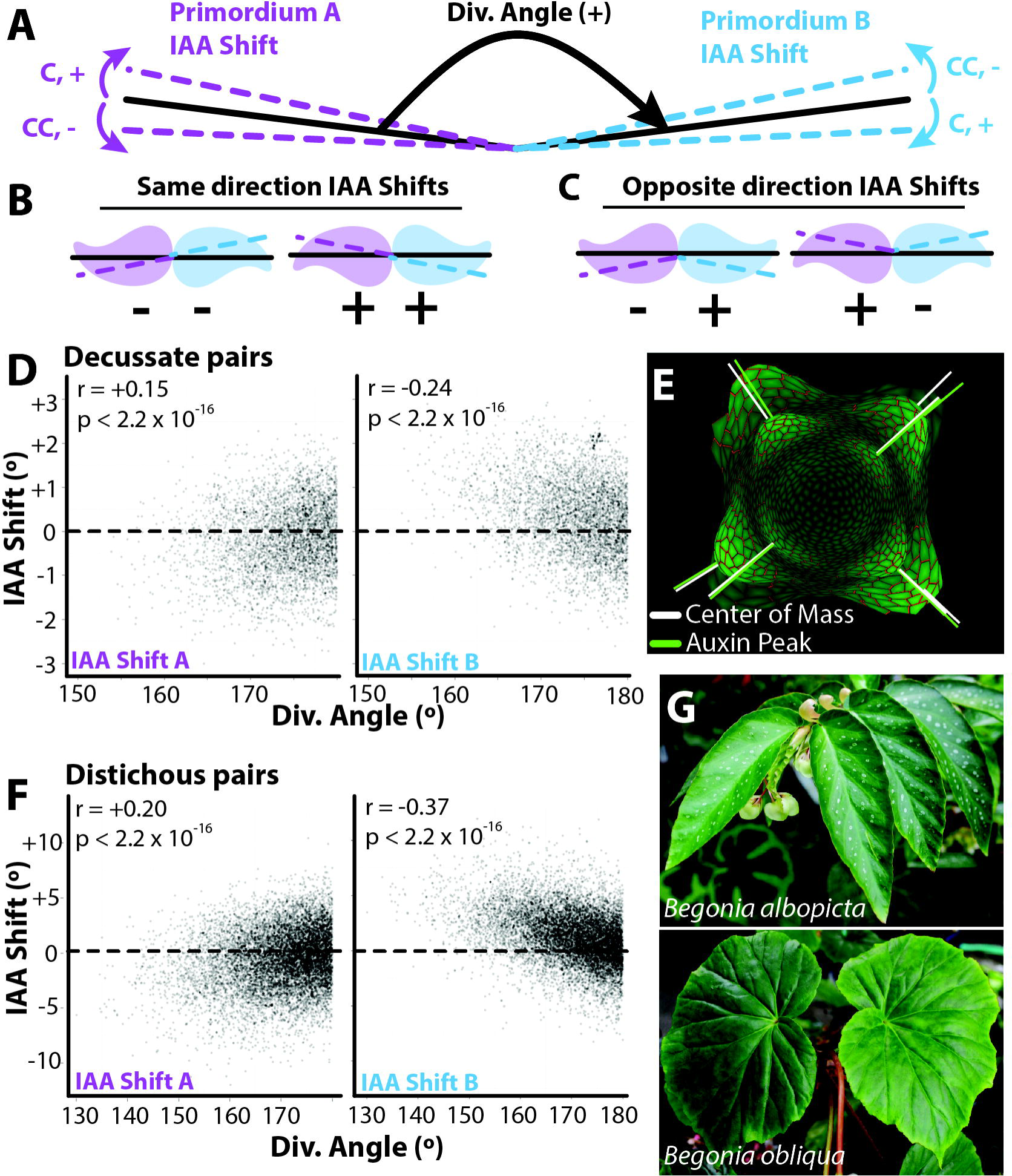
Models of decussate and distichous phyllotaxis predict reflection within leaf pairs. A) Pairs of leaves from decussate and distichous phyllotactic simulations were analyzed such that all pairs have positive divergence angles.IAA Shift values in auxin concentration are then analyzed for “primordium A” and “primordium B” as indicated in purple and blue dotted lines, respectively. Sign (+ or ‐) and direction (C, clockwise and CC, counter-clockwise) of IAA Shift values are as indicated. **B-C**) Diagrams demonstrating the relative mirrored relationships between primordia pairs if IAA Shift values share the same sign and direction (B, “pinwheel,” ++ or ‐‐) or are opposite in sign (C, “mirror,” +‐ or ‐+). **D-F**) As divergence angles deviate from 180° and become smaller, auxin peak concentrations remain truer to an idealized 180° angle. As the divergence angle decreases, primordium A IAA Shift values decrease, whereas IAA Shift values increase under similar conditions for primordium B, such that the discrepancy between auxin and primordia center of masses are mirrored between the two primordia. Results from **D**) decussate and **F**) distichous models are shown, as well as **E**) an image from the decussate model showing primordia (white) and auxin (green) center of masses. **G**) *Begonia* species are excellent examples of reflected, mirrored leaf morphology between distichous leaf pairs, which sometimes manifests as a spiral morphology (lower panel).

### Decussate *e-2* tomato plants display mirrored asymmetries in leaflet position but not terminal leaflet shape

We investigated if auxin asymmetries observed in the decussate model had empirical consequences for leaf morphology in a tomato mutant. Ideally, hypotheses regarding the effects of different phyllotactic systems would not be made between disparate species (such as tomato and *Begonia*) but within a single species. Although mutations that create aberrant phyllotaxis are commonplace, those that produce true transformations from one stable phyllotactic patterning form to another are exceedingly rare [19]. We took advantage of the *e-2* mutation, which has been shown to display decussate-like phyllotaxis, where the divergence angle between leaves 1 and 2 and leaves 3 and 4 approaches 180°, but the angle between leaves 2 and 3 is much smaller (Figure 3A-B) [16].

**Figure 3:**
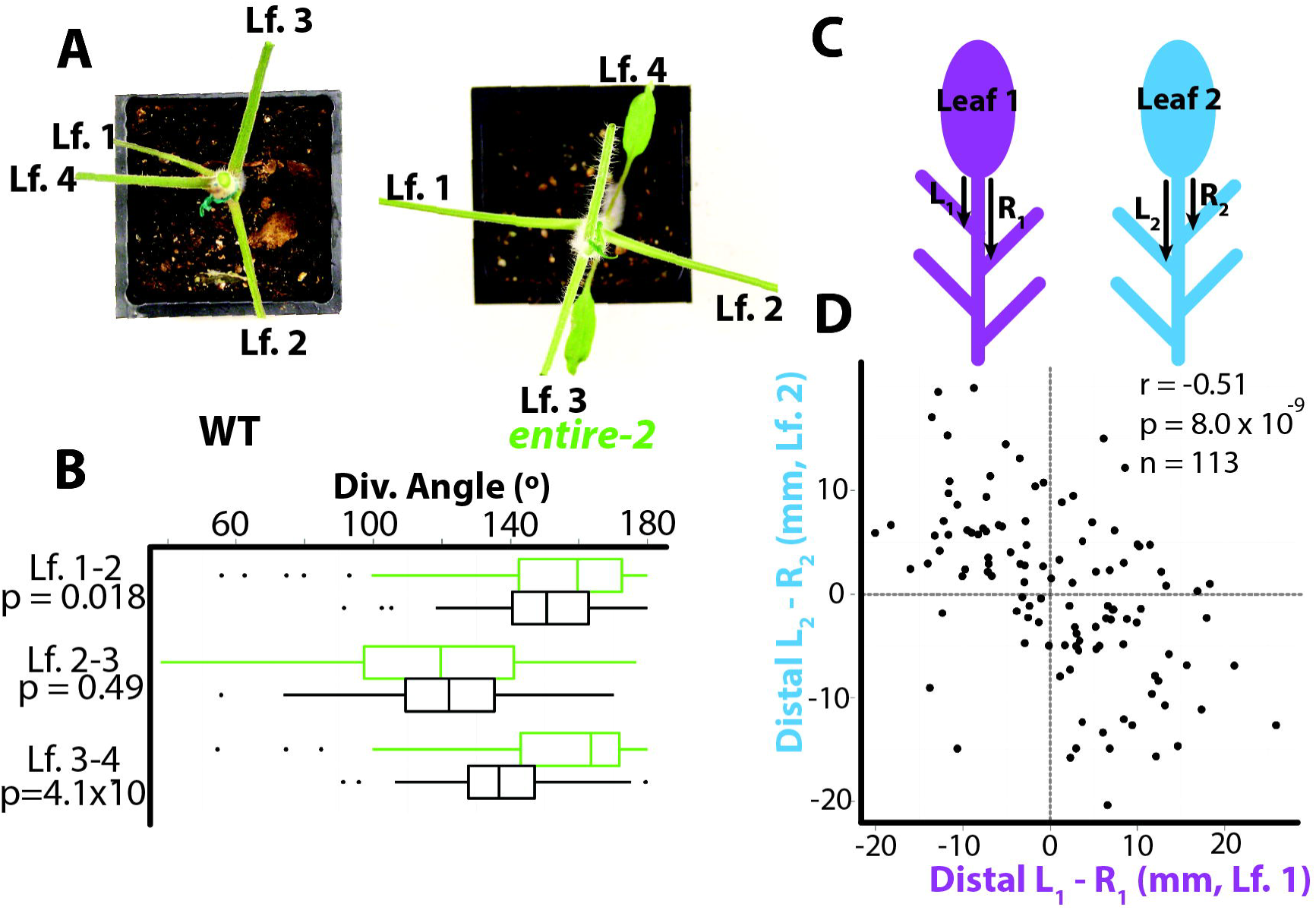
The decussate-like mutant *e-2*. **A)** Relative to a WT background genotype, *entire-2* (*e-2*) exhibits decussate-like phyllotactic patterning. Leaflets have been removed to better view divergence angles. B) Between leaf pairs 1/2 and 3/4 *e-2* divergence angles more closely approximate 180° compared to background genotype. C) “L” and “R” depict the lengths measured to analyze shifts in lateral leaflet positioning on leaf 1 (L_1_ – R_1_) and leaf 2 (L_2_ – R_2_). D) Lateral leaflets on the left and right sides of leaves show a significant negative correlation in the lateral leaflet positioning between leaves 1 and 2, indicative of a mirrored relationship between leaf pairs.

In our previous study [5], the displacement of leaflet position between the left and right sides of the tomato compound leaf was a strong indicator of asymmetry and predicted auxin distributions from the spiral phyllotactic model. For those decussate pairs formed from the first two leaves for which there were leaflets to measure (n=113), we measured the difference in distance from the terminal leaflet to the first distal lateral leaflet between the left and right side of *e-2* leaves (Figure 3C). The direction of the asymmetry of the displacement of leaflets was strongly opposite and mirrored between leaves 1 and 2 in *e-2*, which form a divergence angle greater than that found in wild-type and closer to 180° (Figure 3B). The correlation between the left-right displacement in leaves 1 and 2 in *e-2* is strongly negative (r = ‐0.51, p = 8 × 10^−9^, n=113) (Figure 3D), confirming the mirrored relationship between pairs predicted by modeling (Figure 2D), and distinctly different from the leaflet displacement in the same direction common to leaves arising from wild-type tomato plants with spiral phyllotaxy as we have shown previously [5].

In our previous work on spiral phyllotaxy, we had analyzed terminal leaflet shapes using Elliptical Fourier Descriptors (EFD) and found that leaves arising from plants of a particular handedness were asymmetric in a similar direction. Analyzing leaf1/leaf2 pairs in *e-2*, we asked whether an opposite orientation in asymmetric leaflet shape occurred more frequently than a similar orientation by chance (Figure 4A), as predicted by the “mirrored” orientation observed in our model (Figure 2D). To do so, we isolated the asymmetric sources of EFD variance (Figure 4B) and asked how often each resulting Principal Component (PC) was of the opposite or same sign between leaf1/leaf2 pairs in *e-2*. A one-tailed Fisher’s exact test (in the direction of the alternative hypothesis of more opposite pairs relative to same, assuming equal numbers) fails to achieve significance, leaving the modeled hypothesis of mirrored asymmetry unconfirmed with respect to terminal leaflet shape (Table 1).

**Figure 4:**
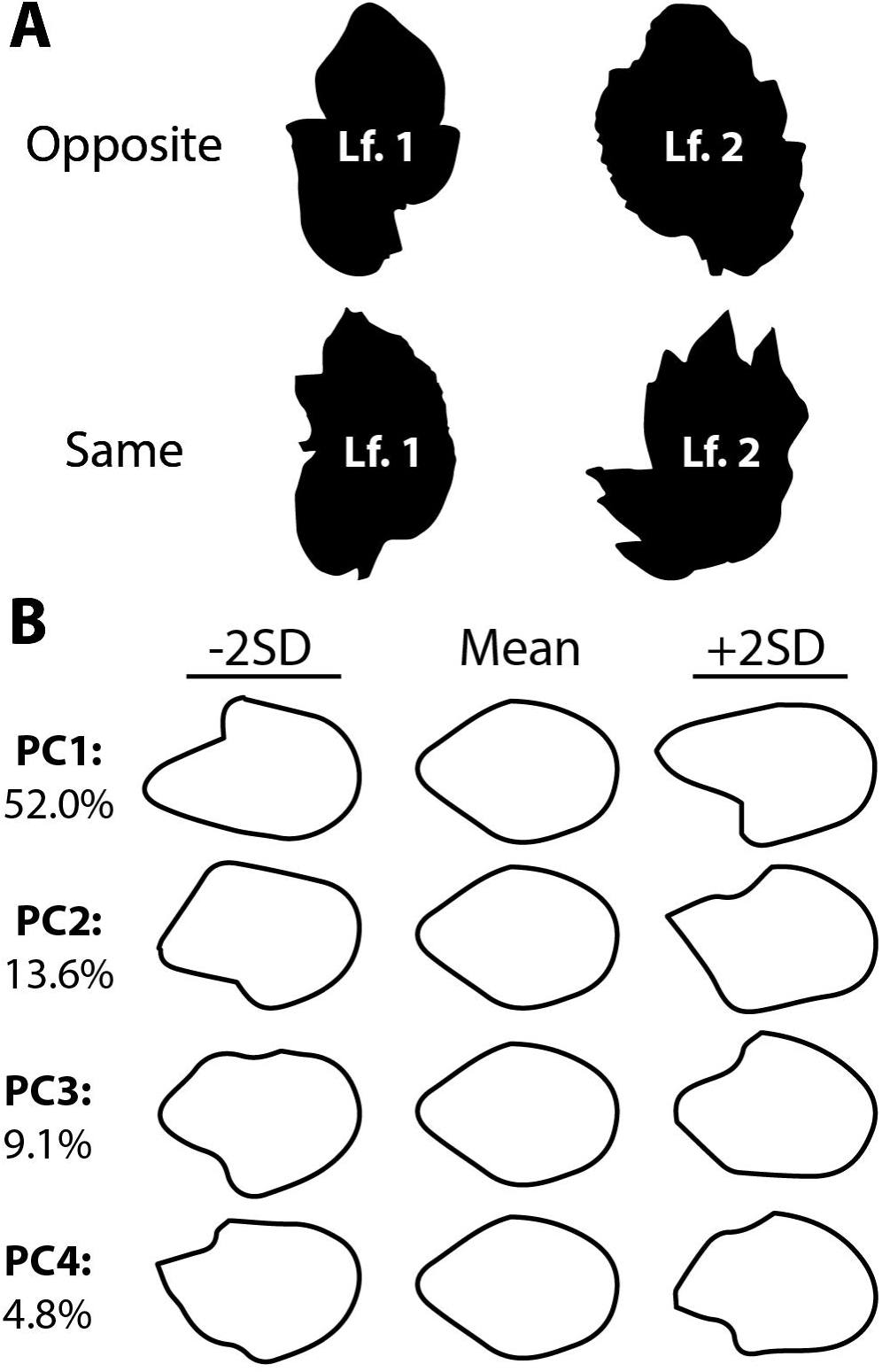
Morphometric analysis of *e-2* terminal leaflets. A) Distal bulges in terminal leaflet laminae in the *e-2* mutant background occur either on opposite sides of leaf 1/2 pairs or on the same side.B) Principal Components (PCs) representing asymmetric shape variance using Elliptical Fourier Descriptors (EFDs). Negative PC standard deviation values are of opposite asymmetric orientation compared to positive values. Percent asymmetric shape variance explained by each PC is provided.

**Table 1:** Opposite vs. same orientation in *e-2* leaf1/leaf2 pairs. For each PC representing asymmetric shape variance in the terminal leaflets of *e*-2 (**Figure 4B**), the number of leaflet pairs with opposite vs. same PC sign, the number of leaf pairs, the ratio of the number of opposite-to-same pairs, and the p-value for the one-sided exact test that the odds ratio of opposite-to-same pairs exceeds 1 are provided.

| PC | Variance | Opposite | Same | n | Ratio | p value |
| --- | --- | --- | --- | --- | --- | --- |
| PC1 | 52.0% | 89 | 94 | 183 | 0.95 | 0.64 |
| PC2 | 13.6% | 87 | 96 | 183 | 0.91 | 0.72 |
| PC3 | 9.1% | 80 | 103 | 183 | 0.78 | 0.91 |
| PC4 | 4.8% | 89 | 94 | 183 | 0.95 | 0.64 |

The mirrored relationship in terminal leaflet pairs may have failed to achieve significance for any number of reasons, including a subtle effect not detected because of insufficient replication (n = 183, which is small compared to the replication measured for common ivy or grapevine species, see below) or that directionalized asymmetry may only be observed in leaflet displacement (Figure 3D) and not terminal leaflet shape. This last hypothesis is intriguing, considering that for all PCs, more same orientations were observed than opposite (Table 1). Considering that tomato is a spiral phyllotactic species and that this is a mutant, perhaps the spiral-phyllotactic tendency for terminal leaflet asymmetries to be of the same sign [5] is preserved in an *e-2* genetic background. Regardless, the mirrored displacements in leaflet position (Figure 3C-D) are predictive of the modeled mirror relationship (Figure 2D) and distinct from what we had previously observed in leaflet position in a wild type background [5], suggesting that some leaf features are affected more by the *e-2* decussate transformation than others.

### Alternating leaf asymmetry in the distichous phyllotaxy of common ivy (*Hedera helix*) and grapevine (*Vitis* and *Ampelopsis* spp.)

To test whether the modeled mirrored predictions for distichous phyllotaxy (Figure 2F) are empirically observable, we turned to distichous vine species, which present numerous numbers of paired, successive nodes to test our hypotheses.

We analyzed 824 leaf pairs of a single genotype of common ivy (*Hedera helix*) growing up a >6m concrete wall enclosing a courtyard at the Donald Danforth Plant Science Center in St. Louis, MO (Figure 5A). Asymmetric shape variance was quantified using Elliptical Fourier Descriptors (EFDs) (Figure 5B). For the first four asymmetric Principal Components (PCs) explaining 87.5% of asymmetric shape variance, more opposite than same signed relationships were detected between leaf pairs (Table 2). However, only for PCs 2 and 3 was the occurrence of opposite-signed leaf pairs significantly greater than that of same-signed pairs at a p value less than the *α* level of 0.05 using a one-way Fisher’s exact test (Table 2). We conclude that especially for specific asymmetric leaf features (defined by PCs 2 and 3 in this particular case) the distichous phyllotactic condition in common ivy confers an alternating, mirrored asymmetry to successive leaves, as predicted by modeling (Figure 2F).

**Figure 5:**
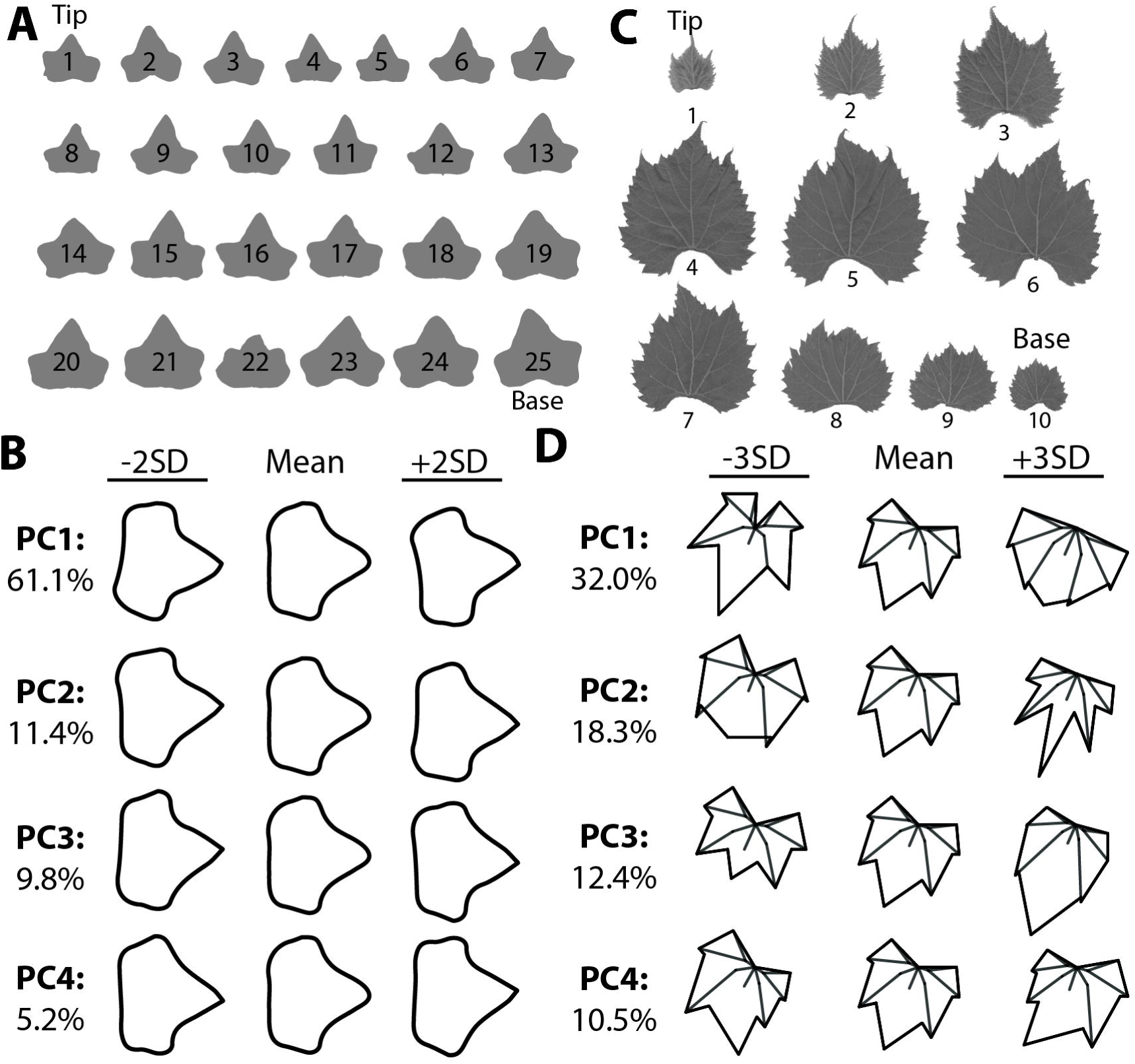
Morphometric analysis of ivy and grapevine leaves. A) An example of leaf outlines analyzed from a shoot of common ivy (*Hedera helix*).Shoot tip and base are indicated. B) Principal Components (PCs) representing asymmetric shape variance using Elliptical Fourier Descriptors (EFDs). Negative PC standard deviation values are of opposite asymmetric orientation compared to positive values. Percent asymmetric shape variance explained by each PC is provided. C) An example of leaf outlines analyzed from a grapevine shoot (specifically, *Vitis xdoaniana*, for which each of the nine leaf pairs have an opposite orientation, see **Table 4**). The analyzable leaf pairs for a shoot equals n-1, where n = total leaf number. In this study, 20 different *Vitis* and *Ampelopsis* (a closely-related genus to *Vitis*) species and hybrids were analyzed. D) PCs representing shape variance in 17 Procrustes-adjusted homologous landmarks. Note that almost all asymmetric shape variance is restricted to PC4, explaining 10.5% of total shape variance, which is analyzed in **Table 4**.

**Table 2:** Opposite vs. same orientation in common ivy (*Hedera helix*) leaf pairs. For each PC representing asymmetric shape variance in ivy leaves (Figure 5B), the number of leaf pairs with opposite vs. same PC sign, the number of leaf pairs, the ratio of the number of opposite-to-same pairs, and the p-value for the one-sided exact test that the odds ratio of opposite-to-same pairs exceeds 1 are provided.

| PC | Variance | Opposite | Same | n | Ratio | p value |
| --- | --- | --- | --- | --- | --- | --- |
| PC1 | 61.1% | 430 | 394 | 824 | 1.09 | 0.2011 |
| PC2 | 11.4% | 462 | 362 | 824 | 1.28 | 0.0078 |
| PC3 | 9.8% | 451 | 373 | 824 | 1.21 | 0.0304 |
| PC4 | 5.2% | 437 | 387 | 824 | 1.13 | 0.1184 |

Next, we analyzed >3,000 leaf pairs from 20 different *Vitis* and *Ampelopsis* species and hybrids from a germplasm collection maintained by the USDA in Geneva, NY (USA). This previously published dataset [20] analyzed leaf shape in >270 vines by collecting all the leaves from a single shoot and recording their order (Figure 5C). Using 17 homologous landmarks superimposed using a Procrustes analysis, both symmetric and asymmetric sources of shape variance are analyzed, but asymmetry is almost exclusively restricted to a specific Principal Component (PC), in this case PC4 which explains 10.5% of the total shape variance (Figure 5D). Performing a one-way Fisher’s exact test to determine if successive leaf pairs with opposite PC4 signs are more prevalent than same-signed pairs, reveals a strong enrichment of mirrored, asymmetric leaf pairs across numerous *Vitis* and *Ampelopsis* species and some hybrids (Table 3). Our results demonstrate that although not necessarily apparent at first glance, grapevine and related species exhibit strong alternating asymmetries in successive leaves (Figure 5C-D; Table 3) consistent with the predictions from models of auxin flux in distichous systems (Figure 2F).

**Table 3:**
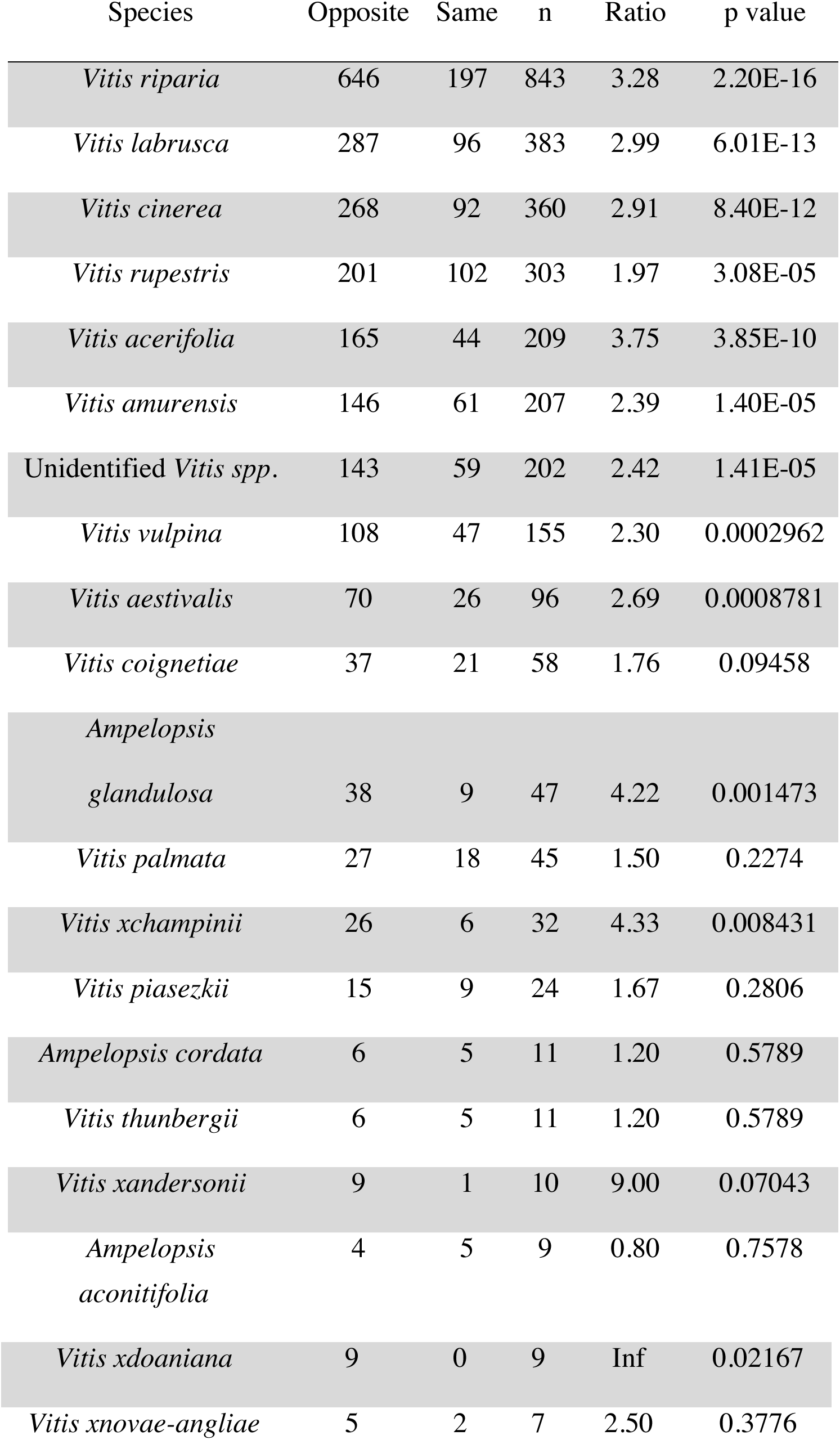
Opposite vs. same orientation in grapevine (*Vitis* and *Ampelopsis* spp.) leaf pairs. For different *Vitis* and *Ampelopsis* species, and *Vitis* hybrids, the number of leaf pairs with opposite vs. same PC4 sign (Figure 5D), the number of leaf pairs, the ratio of the number of opposite-to-same pairs, and the p-value for the one-sided exact test that the odds ratio of opposite-to-same pairs exceeds 1 are provided.

## DISCUSSION

Although they are strikingly different patterning events, our results elucidate common themes between the asymmetries that arise in spiral, decussate, and distichous phyllotaxis (Figure 6). In spiral systems, the peak auxin concentration is displaced towards the descending side of the center of mass of leaf primordia, resulting in a distal shift of morphological features (including leaflet placement and terminal leaflet shape) in mature leaves. The distal shift of morphological features is observed empirically in leaf primordia that lunge towards the ascending side due to increased growth along their descending edge (Figure 6A) [5], reminiscent of the curvatures seen in more extremely asymmetric leaved species, such as leaf shape in *Calathea* or the positioning of serrations in holly [7, 21]. A similar scenario occurs in decussate and distichous systems, except that deviation in divergence angles creates angles less than 180° that, when accompanied by shifts in auxin peak concentrations that remain truer to 180°, creates mirrored morphologies in leaf pairs (Figure 2F; Figure 6B-D). The mirrored relationship between leaf pairs is strikingly observed in the left-right displacement of leaflets in the leaves of the decussate tomato mutant *entire-2* (Figure 3) but not in the shape of the terminal leaflets (Figure 4; Table 1), perhaps reflecting that tomato is inherently spirally phyllotactic and not all leaf features reflect the transition to decussate phyllotaxy this mutation confers. Two different vine species, common ivy and grapevine and its relatives, are statistically over-represented for opposite-signed leaf asymmetries compared to same-signed asymmetries (Figure 5; Tables 2-3), grapevine particularly so, again confirming the mirrored relationship predicted in the distichous model (Figure 2F).

**Figure 6:**
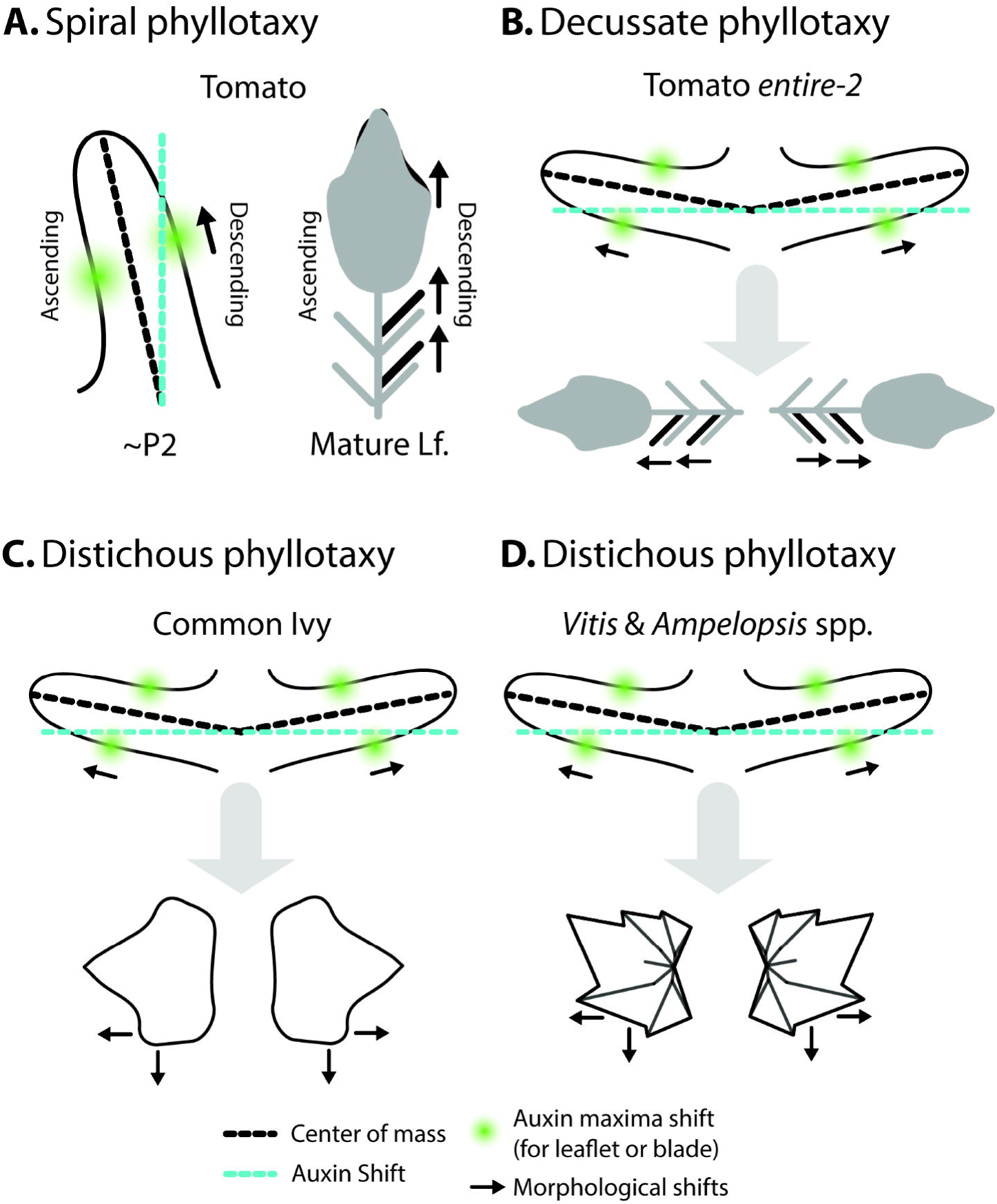
Summary of leaf asymmetry in spiral, decussate, and distichous systems. The discrepancy between the peak concentration of auxin (blue) and the respective primordium center of mass (black) is such that the auxin falls towards a more idealized phyllotactic position.In both A) spiral and B-D) decussate/distichous systems, this results in the distal shifting of morphological features (such as leaflets, terminal leaflet lobes) and increase laminar outgrowth on the side of the primordium with greater auxin concentration. Asymmetries early in leaf development manifest as the shifting of distal features on the descending side of mature leaves in spiral systems and mirroring of leaf features in decussate/distichous systems.

Although we have focused on leaf shape, it is important to remember that auxin potentially impacts many more features of the leaf than just shape, or that leaf shape is a read-out of other underlying asymmetries at the cellular and tissue levels [15, 22, 23]. Further, it is possible that the patterning of auxin itself is influenced by properties specific to particular shoot apical meristem architectures. In this regard, it is important that the underlying developmental causes of leaf asymmetry be examined further. For example, the archetypal left-right leaf asymmetry of extreme midrib and blade curvature and spiraling seen in distichous *Begonia spp*. leaf pairs (Figure 2G) is apparent in very early primordia [24, 25]. In fact, in *Begonia* mirrored asymmetry appears to be the null condition, and symmetric *Begonia* have asymmetric primordia that become symmetric subsequently during development [24]. If leaf primordia are inherently asymmetric and symmetry is a result of secondary development, it is perhaps not so surprising that morphometric methods can statistically detect directional asymmetries in superficially symmetric species. We also note that the shoot apical meristem of grapevine and related species is dorsiventrally patterned perpendicular to the axis of distichous leaf initiation [26]. Different numbers of vascular traces enter the leaves on the dorsal and ventral sides of the shoot, which in the context of distichous phyllotaxy, creates an inherent mirrored asymmetry in vascular patterning, perhaps explaining the extremely strong mirrored asymmetry we observe in grapevine (Table 3).

Different phyllotactic systems—spiral, decussate, and distichous, among others—create inherently different spatial relationships between leaf primordia in the shoot apical meristem.The flux of auxin between leaf primordia especially accentuates differences along the left-right axis. Considering the global roles that auxin plays in vascular patterning and leaf shape, it is impressive that most leaves appear as bilaterally symmetric as they do. Our results clearly demonstrate that left-right leaf asymmetry, reflecting predicted auxin asymmetries in the shoot apical meristem, is influenced by phyllotaxy and present in diverse species.

## MATERIALS AND METHODS

This work was conducted in parallel and together with our work on spiral phyllotaxy [5]. Some of the materials and methods from Chitwood, Headland et al., 2012 and Chitwood et al., 2016 are repeated here for convenience.

### Plant Material and Growth and Conditions

*Solanum lycopersicum* accession LA3475 (cv. M82) was used for “wild-type” tomato measurements. The *entire-2* accession used for this study is 3-705. Tomato resources were obtained from the U.C. Davis Tomato Genetics Resource Center (TGRC). Tomato seed was sterilized for 2 minutes in 50% bleach, washed in water, and plated onto wet paper towels in Phytatrays (Sigma). Seed was kept at room temperature in darkness for 3 days and then transferred to chamber conditions in light for an additional three days before transplanting into Sunshine soil mix (Sun-Gro Horiculture). For measures of leaflet positioning and terminal leaflet shape, tomato plants were analyzed 33 days after plating seed.

Common ivy leaves (*Hedera helix*) were obtained from a single genotype of vine growing up >6m walls enclosing a courtyard at the Donald Danforth Plant Science Center in St. Louis, MO December 28, 2015. Documentation of the site location and material collected can be found in the following link (https://twitter.com/DanChitwood/status/681554095618965504). The leaves from 41 vines, with up to 25 leaves each, were dissected from the shoot and scanned.

Leaves from 20 different grapevine species and hybrids (*Vitis* and *Ampelopsis* spp.) were collected from >270 vines in the USDA *Vitis* germplasm collection in Geneva, NY in June 2013 [20]. A single shoot was sampled from each vine, the leaves dissected, scanned, and their position in the shoot noted.

### Measures of lateral leaflet displacement and asymmetric shape

Measurements of lateral leaflet position and terminal leaflet shape in tomato were made from photographs. The first four leaves of tomato plants were dissected, placed under non-reflective glass, and their terminal leaflets removed at the base. Photos of the leaf series were taken using Olympus SP-500 UZ cameras mounted on copy stands (Adorama, 36’’ Deluxe Copy Stand) and controlled remotely by computer using Cam2Com software (Sabsik). Leaves from ivy and grapevine were dissected from the shoot and scanned (Epson Workforce DS-50000, Suwa, Japan). Care was taken to include the node position of each leaf in the scan.

In tomato, measures of the distance from the base of the terminal leaflet or the petiole to the most distal and proximal leaflets were made using measurement functions in ImageJ. Lengths were normalized using rulers present in each photograph. For shape analysis in tomato and ivy, photographs were first converted to binary form using ImageJ and individual leaflets extracted from the leaf series and named appropriately as separate files.

The analysis of tomato and ivy leaflet and leaf shape (respectively) was conducted using Elliptical Fourier Descriptors followed by PCA using the program SHAPE [27]. Object contours were extracted as chain-code. Chain-code was subsequently used to calculate normalized EFDs. Normalization was based upon manual orientation with respect to the proximal-distal axis of the leaflet/leaf. Principal component analysis was performed on the EFDs resulting from the first 20 harmonics of Fourier coefficients. Only asymmetric sources of shape variance were analyzed using the *b* and *c* coefficients [28]. Coefficients of Elliptical Fourier Descriptors were calculated at −2 and +2 standard deviations for each principal component and the respective contour shapes reconstructed from an inverse Fourier transformation. PCs were then subsequently analyzed.

For grapevine data, 17 landmarks were placed, in order, for each leaf using the ImageJ [29] point tool. Landmarks and their order were as follows: 1) petiolar junction, 2) midvein tip, 3) left distal sinus, 4) right distal sinus, 5) left distal lobe tip, 6) right distal lobe tip, 7) left proximal sinus, 8) right proximal sinus, 9) left proximal lobe tip, 10) right proximal lobe tip, 11) left terminus petiolar vein, 12) right terminus petiolar vein, 13) branch point midvein, 14) branch point left distal vein, 15) branch point right distal vein, 16) branch point left proximal vein, 17) branch point right proximal vein (see Chitwood et al., 2016 for a visualization of landmark position). Using ggplot2 [30] in R [31], graphs for landmarks from each image were visually checked for errors. If errors were detected, the landmarking was redone for those particular samples. Once a quality landmarked dataset was created, a Generalized Procrustes Analysis (GPA) was undertaken using the R package shapes [32]. Eigenleaves were visualized using the shapepca function and PC scores, percent variance explained by each PC, and Procrustes-adjusted coordinates were obtained from procGPA object values.

#### Statistical analysis

All basic statistic functions were performed in R [31] and visualized in the package ggplot2 [30]. The cor.test() function in the stats package was used for all correlation analyses using method = “pearson”. Fisher’s exact test was performed using fisher.test() and the one-way test used to determine if the odds ratio of opposite-to-same orientations was greater than 1 by setting alternative = “greater”.

#### Auxin transport modeling

The simulation models have been described previously [5], however we summarize here for completeness. Cells are modelled as polygons on a growing apex surface and thus are assumed to be uniform in thickness. Extracellular space is ignored, and diffusion and transport occur directly from cell to cell. The change in concentration of auxin in a cell is modeled as:

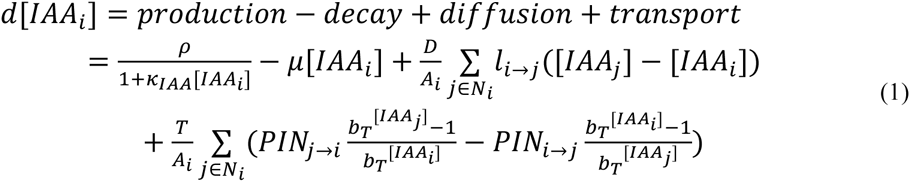

where [*lAA*_*i*_] is the concentration of auxin in cell *i*, *ρ* controls the rate of production with saturation coefficient *k*_*IAA*_, *μ* controls decay, *D* is the diffusion coefficient, *A*_*i*_ is the area of cell *i*, *N*_*i*_ are the neighbours of cell *i*, *l*_*i*→*j*_ is the length of the wall between cell *i* and *j*, *T* is the transport coefficient, *PIN*_*i*→*j*_ is the amount of PIN on the membrane of cell *i* facing cell *j*, and *b*_*T*_ is the base for exponential transport. Equation (1) was used for the decussate simulation, but in the distichous simulation we used the original quadratic transport term from Smith et al. (2006) [13]:

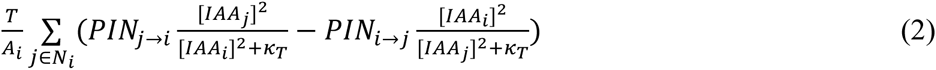

where *κ*_*T*_ controls saturation of auxin transport. We model PIN allocation to the membranes as:

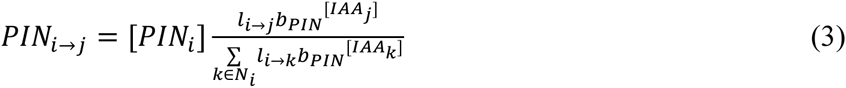

where [*PIN*_*i*_] is the total amount of PIN in cell *i*, and *b*_*PIN*_ is the base for exponential PIN allocation to cell membrane sections. Equations (1-3) where implemented on the growing cellular template as described in Smith et al. (2006) [13]. Simulations were performed using Vertex-Vertex systems [33] in the Vlab modeling environment [34].

## ACKNOWLEDGEMENTS

C.C.M was supported by a National Science Foundation Graduate Research Fellowship (DGE-1148897), Katherine Esau Summer Fellowship, Walter R. and Roselinde H. Russell Fellowship, and Elsie Taylor Stocking Fellowship. Part of the work was supported by NSF PGRP grant IOS–0820854 (to N.R.S., Julin Maloof and Jie Peng). Funding for R.S.S. was provided by the SystemsX.ch Plant Growth Research, Technology, and Development project and the Max Planck Society.

## AUTHOR CONTRIBUTIONS

C.C.M, D.H.C, R.S.S, & N.R.S designed the experiments, performed the experiments, analyzed the data and wrote the paper.

